# Modeling Temporal Dependencies and Feature Interactions Reveal Novel Clinical and Molecular Insights into Alzheimer’s Disease Progression

**DOI:** 10.1101/2025.11.27.690894

**Authors:** A H M Osama Haque, Abdullah Al Fahad, M Sohel Rahman, Md Abul Hassan Samee

**Affiliations:** Department of Computer Science and Engineering, Bangladesh Univeristy of Engineering and Technology, Dhaka, 1000, Bangladesh; Department of Computer Science and Engineering, United International University, Madani Avenue, Dhaka-1212, Bangladesh; Department of Computer Science, Indiana University, 835 paca st, Indianapolis, 46202, Indiana, IN, United States of America; Integrated Physiology, Baylor College of Medicine, Houston, TX 77030, United States of America

**Keywords:** Interpretability, Feature interaction, TreeSHAP, Integrated Gradient, ADNI, TADPOLE, Temporal dynamics

## Abstract

**Objective:** Alzheimer’s Disease remains a major public health challenge, requiring insights into feature interactions and temporal trends of feature importance. Community-wide data science competitions such as the TADPOLE Challenge provide platforms to benchmark predictive models using ADNI datasets. While top-performing models achieve accurate predictions, they often leave mechanistic questions unresolved. We introduce a framework that separately models static feature interactions and temporal dynamics, enabling complementary insights from longitudinal AD data.

**Materials and Methods:** We analyzed XGBoost with TreeSHAP for feature interactions and RNN-AD with Integrated Gradients for temporal trends. This two-branch design allows XGBoost to capture nonlinear cross-sectional interactions, while RNN captures evolving, time-dependent influences. The attributions are fused into a combined importance map.

**Results:** Our framework showed agreement between TreeSHAP and IG, highlighting FAQ, CDRSB, ADAS13, MMSE, and RAVLT variants as the most consistently important features across both branches. Temporal attribution analysis revealed stage-dependent trends: in CN, features such as DX:CN, FAQ, and RAVLT_immediate increased in importance with longer prediction horizons; in MCI, MidTemp and WholeBrain gained importance; and in AD, FAQ remained dominant. Feature-interaction analysis identified strong clinical–clinical interactions and secondary clinical–molecular interactions involving hippocampal and entorhinal volumes.

**Discussion:** Combining interaction and temporal trends showed that RAVLT_immediate, FAQ, and DX-based features were the only markers consistently influential across both dimensions, indicating stable, cross-validated predictors of Alzheimer’s disease progression. Feature importance in AD prediction is dynamic, with early-time features often most influential. These insights support personalized monitoring, adaptive modeling, and mechanistic interpretability, enhancing patient-specific interventions and trial design.

**Conclusion:** This work highlights feature interactions and temporal trends in AD prediction models, offering insights for personalized treatments and patient-specific trial designs. Our framework provides stable, cross-validated explanations that unify structural and temporal importance, enhancing trustworthiness and mechanistic interpretability in AD modeling.

## INTRODUCTION

Alzheimer’s disease (AD), characterized by progressive neurodegeneration and cognitive decline, represents one of the most pressing healthcare challenges as global populations age ^1–3^. Machine learning models have achieved remarkable success in AD prediction, leveraging multimodal data, including clinical histories, cognitive assessments, neuroimaging, and genetic biomarkers, to enable early diagnosis and prognosis ^4–7^. Accurate and early diagnosis is pivotal for the success of both protective and preventive treatment strategies, yet diagnosis remains challenging and often subjective, with sensitivity and specificity ranges indicating risks of misclassification ^8^. Furthermore, predicting disease progression poses additional challenges, motivating the use of computational and AI-based systems for more reliable prognostication ^9,10^.

The TADPOLE challenge exemplified this progress, with models like MinimalRNN (CBIL) and XGBoost (Frog) demonstrating exceptional predictive performance for individual AD trajectories ^11–14^. Subsequent studies have further refined these approaches, employing feature selection ^15^, CNN architectures ^16^, and deep RNNs ^17^ to enhance prediction accuracy.

Despite these predictive successes, a critical gap remains: these models operate as “black boxes,” providing predictions without offering mechanistic insights into disease progression. While recent work has begun applying explainable AI (XAI) techniques to AD models— with studies using SHAP, LIME, and GradCAM to assess feature contributions ^18,19^—there remains significant room for improvement. Hernandez et al. ^20^ and El-Sappagh et al. ^21^ utilized SHAP for feature importance analysis; however, neither explored how features interact with each other nor how their importance evolves. This limitation is particularly problematic for AD, a progressive disease where biomarker relevance shifts across disease stages and prediction horizons.

Understanding these dynamics is essential for AD specifically because the disease’s temporal nature means that different biological processes dominate at different stages. Molecular changes may characterize early stages, while later stages show pronounced clinical manifestations. Without understanding feature interactions, how clinical markers influence molecular ones and vice versa, and temporal trends in feature importance, we miss opportunities to develop stage-specific interventions and time-sensitive treatment strategies.

To address these challenges, we propose an interpretability-driven, dual-branch framework tailored for longitudinal AD data. Longitudinal datasets present two inherent complexities: (i) temporal dynamics evolving across multiple time points, and (ii) complex feature interactions occurring within each time point. Rather than relying on a single architecture to capture both behaviors, our approach employs two complementary models in parallel, one optimized for temporal reasoning and the other for static feature interactions.

This study addresses these critical gaps through two novel contributions. First, we perform the first comprehensive analysis of feature interactions in AD prediction, systematically examining clinical–clinical, molecular– molecular, and clinical–molecular feature pairs in XG-Boost models. Our analysis reveals, for instance, that the interaction between FAQ scores and ABETA levels provides stronger predictive power than either feature alone, suggesting compensatory mechanisms between functional decline and amyloid burden. Second, we present the first detailed temporal interpretation of RNN-based AD models, uncovering how feature importance evolves with prediction horizon. We demonstrate that for cognitively normal (CN) subjects, clinical features such as DX:CN and FAQ show steadily increasing importance from 1-month to 6-month predictions, whereas for mild cognitive impairment (MCI) subjects, molecular features like MidTemp and WholeBrain volume exhibit similar temporal trends. These findings suggest that optimal biomarker selection should align not only with disease stage but also with the intended prediction timeframe.

By combining two complementary interpretability branches, our framework produces explanations that are more stable, robust, and trustworthy than single-method attribution approaches, an important requirement for clinical decision-support and biomarker discovery. By moving beyond static feature importance to examine interactions and temporal dynamics, our work transforms black-box AD models into interpretable systems that reveal disease mechanisms. This interpretability is crucial for developing targeted interventions, validating biological hypotheses, and ultimately improving patient outcomes in this devastating disease.

Moreover, the framework we present holds broad applicability beyond AD. Other progressive diseases with temporal and multimodal complexity, such as frontotemporal dementia, Parkinson’s disease, and atrial fibrillation, stand to benefit from similar approaches to feature interaction analysis and temporal attribution.

## MATERIALS AND METHODS

### Dataset

TADPOLE refers to “The Alzheimer’s Disease Prediction Of Longitudinal Evolution”. TADPOLE challenge aims to predict the early symptoms reflected in people with a high risk of AD, preferably in short to medium term (1-5 years). Alzheimer’s disease neuroimaging initiative (ADNI) study provides longitudinal data to help investigate research questions regarding the progression of dementia, data specificity and models to best forecast future measurements. The TADPOLE challenge comes with three standard datasets (1677 patients) that are acquired from the ADNI study.

#### Notable Biomarkers

Among the main cognitive tests, there are CDRSumof-Boxes(CDRSB), ADAS11, ADAS13, MMSE, RAVLT, Moca, etc. Rey’s Auditory Verbal Learning Test (RAVLT) is a widely used neuropsychological assessment tool designed to evaluate episodic memory function ^22^. It serves as an effective measure in dementia and pre-dementia evaluations, with impaired RAVLT scores correlating strongly with Alzheimer’s disease pathology. The Alzheimer’s Disease Assessment Scale–Cognitive Subscale (ADAS-Cog), developed in the 1980s, measures cognitive dysfunction in Alzheimer’s disease ^23^. Its utility in pre-dementia stages has expanded despite concerns about its sensitivity to detect changes at milder disease stages. This review assesses ADAS-Cog’s performance in pre-dementia populations and evaluates 31 modified versions aimed at improving measurement accuracy. Modifications include altering scoring methods and incorporating tests of memory, executive function, and daily functioning, though heterogeneity introduced by these changes may complicate comparisons across studies ^24^.

### Machine Learning Models: RNN and XGBoost

We focused on the Recurrent neural model (RNN-AD) used by team CBIL and the XGBoost model used by team FROG.

#### RNN-AD

Chen et. al. ^25^ proposed minimalRNN, which is incorporated in the RNN-AD architecture defined by Nguyen et. al. for predicting AD disease progression. Minimal-RNN requires fewer parameters than LSTM and therefore, overfitting can be reduced. In this proposed architecture, as depicted in Figure 1 (A),

*x*_*t*_ = All variables observed at time step t

*s*_*t*_ = The diagnosis status

*g*_*t*_ = The continuous values

The MinimalRNN update equations are as follows:

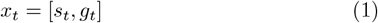

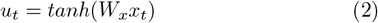

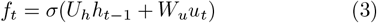

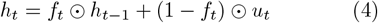

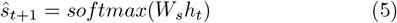

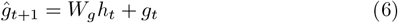

**Figure 1.**
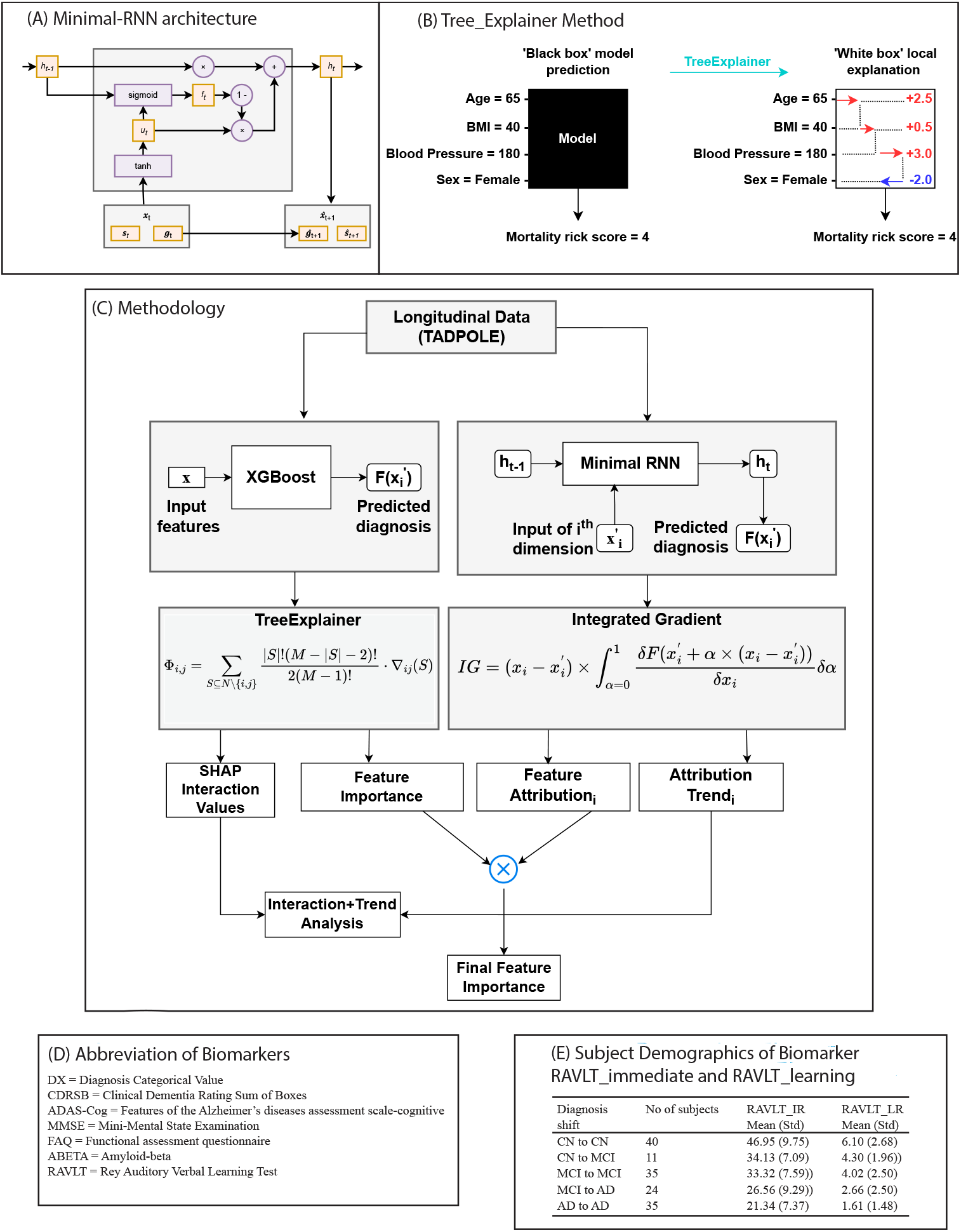
(A) MinimalRNN architecture, (B) TreeSHAP method description, (C) Fusion framework, (D) Biomarkers abbreviation and (E) Subject demographics of RAVLT_immediate and RAVLT_learning to demonstrate their importance

Diagnosis is represented as one-hot encoding of three possible classes, namely Cognitive Normal (CN), Mild Cognitive Impairment (MCI) and AD (Alzheimer’s Disease Dementia). At each time point, hidden state *h*_*t*_ is updated using transformed input *u*_*t*_ and the previous hidden state *h*_*t* − 1_. The hidden state includes information about the previous time steps, which is a significant part of a recurrent neural network. The current hidden state ht is used to predict the observation in the immediate next time point *x*_*t*+1_. The forget gate, denoted as *f*_*t*_, evaluates the influence of both the previous hidden state *h*_*t* − 1_ and the current transformed input *u*_*t*_ on the current hidden state *h*_*t*_ (refer to Equation 3). The model then uses *h*_*t*_ to forecast the diagnosis *s*_*t*+1_ for the next month and continuous variables *g*_*t*+1_ (see Equations 5 and 6. The symbols ⊙ and *σ* represent the element-wise product and the sigmoid function, respectively.

The top-performing system in predicting diagnoses for the TADPOLE Challenge was Frog, which utilized a gradient boosting machine powered by XGBoost ^26^. Gradient boosting is a machine learning technique that builds a powerful predictive model by iteratively combining the predictions of multiple weak learners, such as decision trees, in a way that minimizes the overall prediction error. It achieves this by sequentially fitting new models to the residuals of the previous ones, using gradient descent optimization to minimize a chosen loss function. XGBoost stands for “eXtreme Gradient Boosting.” It’s an optimized and efficient implementation of the gradient boosting algorithm. XGBoost is designed for speed and efficiency, making it capable of handling large datasets with millions of instances and features. It’s highly optimized, using techniques like parallelization and tree pruning to make training and prediction faster. The most important hyperparameters of gradient boosting are the learning rate (learning_rate), maximum tree depth (max_depth), number of trees to fit (n_estimators) and L2 regularization weight (reg_lambda). Hyperparameter selection was performed using a randomized search with RandomizedSearchCV from scikit-learn with five-fold cross-validation and ten iterations. The hyperparameter values were found as learning_rate=0.1 max_depth=8, n_estimators=100, gamma=10.0, reg_lambda: 0.1

### Interpretability

Nguyen et. al. ^27^ proposed minimalRNN for creating the AD progression model because it has fewer parameters than complex LSTM models. Therefore, it is less likely to overfit. In the TADPOLE grand challenge on July 3rd, 2020, their proposed minimalRNN models (labeled as “CBIL-MinMFa” and “CBILMinMF1”) secured 2nd and 3rd place, respectively, in the TADPOLE challenge. Therefore, we opted to interpret this model to better understand the feature importance in predicting biomarkers and diagnosis status using longitudinal data. Regarding recurrent neural networks, conventional explainable AI methods like SHAP are not as effective. Instead, we integrated the “Integrated Gradient” technique from the “Captum” package to emphasize the attribution values of features for each specific class ^28^.

#### Integrated Gradient

Understanding how a model generates predictions and identifying the factors influencing these predictions are crucial aspects of machine learning. Various techniques have been developed for model interpretation, including feature importance analysis, gradient-based methods, and rule-based approaches. Integrated Gradients is a cutting-edge method for model interpretation that adheres to two fundamental principles: sensitivity and implementation invariance. The sensitivity principle asserts that a feature’s contribution to the model’s output should be proportional to its impact on the output, while the implementation invariance principle stipulates that functionally identical models should yield consistent attributions.

Integrated Gradient offers several advantages over other interpretation tools, such as plain gradients, LIME ^29^, DeepLift ^30^, and Layer-wise Relevance Propagation ^31^. Unlike plain gradients, Integrated Gradients consider the entire path between a baseline input and the actual input, providing a more comprehensive understanding of feature importance. It overcomes the limitations of other methods by adhering to both sensitivity and implementation invariance axioms. Wang et al. ^32^ derived heatmaps from CNNs trained on T1 MRI scans from the ADNI dataset and compared these to Support Vector Machine (SVM) activation patterns. Their results showed that Integrated Gradient produced the best overlap result, outperforming SVM patterns and other methods in capturing relevant brain regions. Cik et al. ^33^ also employed methods like Integrated Gradients (IG) and Layer-wise Relevance Propagation (LRP) in their study.

The principle behind Integrated Gradients involves computing the integral of the model’s output gradients with respect to the input, along a straight path from a chosen baseline to the actual input. This integral represents the cumulative effect of each feature along the path and serves as a measure of feature importance. By integrating the gradients over this path, Integrated Gradients generate attribution scores for each feature, indicating their relative contribution to the model’s decision-making process.

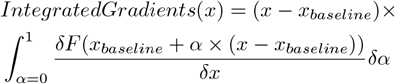

The formula for Integrated Gradients, as presented in the original paper, involves the actual input x, chosen baseline *x*_*baseline*_, model output *F*, and the gradients of the model’s output with respect to the input 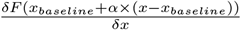. By evaluating this integral for each feature, Integrated Gradients produce attribution scores that quantify the influence of each feature on the model’s predictions. Additionally, Integrated Gradients can be applied to calculate the importance of layers and neurons without necessitating any modifications to the model ^34^.

For the RNN-AD model, we can set a specific duration up to which the predictions are made. Initially, 5 years (5×12 = 60 months) of predictions were made. Using Integrated Gradients, we can generate attribution scores for only a specific duration. Therefore, we conducted results for different durations. For each duration value, we generated attribution scores of the 22 features that contributed to the categorical prediction (CN/MCI/AD) and regression (ADAS11, Ventricles, ICV). When applying integrated gradient, it assumes the model’s forward function to return only a single tensor. However, the original RNN-AD was modified to return a tuple of tensors, one for categorical outputs and the other for regression outputs. Therefore, we ran the simulation set twice: once with the forward function returning categorical outputs and the other returning regression outputs. It is important to note that the model is trained as before, optimizing the loss by integrating both categorical and mean absolute loss.

We focused on the models that performed well according to the analysis of Marinescu et. al. ^14^. We extended the model predictions up to year 5 to better understand the consistency of the feature importance and the effect of the temporal nature of the model.

#### TreeSHAP

SHAP (SHapley Additive exPlanations) ^19^ is a unified framework for interpreting predictions introduced by Lundberg et al. SHAP assigns each feature an importance value for a particular prediction, providing a unique additive feature importance measure that adheres to three key properties: local accuracy, missingness, and consistency. These properties are comprehensively described in the original paper. SHAP values utilize conditional expectations to define simplified inputs. TreeSHAP, introduced by Lundberg et al. ^35^, is a fast method for computing SHAP values specifically for tree-based models. While feature attributions are typically allocated to each input feature, TreeSHAP offers additional insights by separating interaction effects from main effects. A detailed overview of the TreeExplainer method is depicted in Figure 1 (B). Given that SHAP values are based on classic Shapley values from game theory, a natural extension to interaction effects is achieved through the more modern Shapley interaction index:

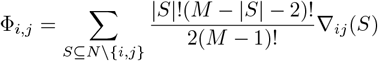

when *i* ≠ *j*, and

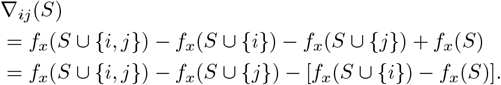

In the Equation, the SHAP interaction value between feature *i* and feature *j* is equally divided between the two features, resulting in *ϕ*_*i,j*_ = *ϕ*_*j,i*_. Consequently, the total interaction effect is given by *ϕ*_*i,j*_ + *ϕ*_*j,i*_. The main effects for a prediction can then be defined as the difference between the SHAP value and the SHAP interaction values for a feature:

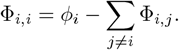

Here *M* is the number of input features, and *ϕ*_*i*_ ∈ ℝ, S is a subset of the input features and *f*_*x*_(*S*) is defined as:

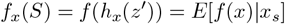

The SHAP library provides various types of visual aids, with the two most important being the interaction plot and the main effect plot.

We adopted the data preprocessing code provided by ssedai026, available at (https://github.com/ssedai026/tadpole-challenge.git). This code was instrumental in preparing our dataset for subsequent analysis. We utilized the D1_D2 Dictionary files and the D1_D2_.csv file to construct our dataset. To facilitate the analysis, we converted the DX column, which represents diagnostic categories, into numerical values: 0 for cognitively normal (CN), 1 for mild cognitive impairment (MCI), and 2 for Alzheimer’s disease (AD). This transformation allowed for more straightforward computational handling and analysis of diagnostic data. In addition to the existing fields, we introduced a new field named ‘Age_r’, which represents the patient’s age in months at each corresponding visit. The calculation of ‘Age_r’ is illustrated in Equation 1, where ‘AGE’ denotes the patient’s age at the baseline (first visit), and ‘month_bl’ indicates the number of months since the baseline visit. Age_r = month_bl + 12 * AGE This new field provides a more granular view of the patient’s age progression over time, which is crucial for our longitudinal analysis. After these preprocessing steps, we extracted the relevant features and labels. These were then grouped on a per-patient basis to ensure that our analysis could account for the individual variability and longitudinal nature of the data. This patient-wise grouping is essential for capturing the temporal dynamics of the disease progression and for making accurate predictions based on each patient’s history. For the test-train split, we adhered to the 80-20 rule, using GroupShuffleSplit to ensure the integrity of groupings within the data. This method helps maintain the distribution of data points, ensuring that the training and testing sets are representative of the overall dataset. In creating our model, we defined an estimator using XGBClassifier with the ‘softmax’ objective function, suitable for multi-class classification tasks. We then defined a search space for the hyperparameters, the values of which are shown in Supplement . Using RandomizedSearchCV, we created a pipeline object that integrated the estimator with a cross-validation splitting strategy, specifically employing five-fold cross-validation. This method allowed us to evaluate the model performance more rigorously across different subsets of the data.

#### Fusion framework

We propose an interpretability-driven framework (Figure 1 (C)) tailored for longitudinal data, which inherently exhibits two major challenges:

1. Temporal dynamics that evolve across multiple timepoints
2. Complex feature interactions that occur within each time point.

Rather than relying on a single architecture to capture both properties, we employ two complementary models in parallel. One trail is designed for temporal reasoning and the other is optimized for static feature interactions. The first branch utilizes XGBoost, a tree-based ensemble method capable of modeling nonlinear interactions while naturally handling missing or irregularly sampled data. XGBoost is trained on aggregated longitudinal features to predict the target clinical outcome. We then apply TreeExplainer (TreeSHAP) to obtain exact Shapley values for each prediction. TreeSHAP quantifies the marginal contribution of each feature and reveals higher-order interactions encoded within the ensemble, providing a comprehensive, global view of structural feature importance across individuals. The second branch employs a Minimal Recurrent Neural Network (RNN), which directly models the sequential nature of the longitudinal trajectories. This network captures temporal dependencies by processing each subject’s time-ordered feature sequence. To interpret this branch, we use Integrated Gradients (IG), which computes feature attributions by integrating gradients along a path from a baseline input to the actual input. IG therefore yields a temporal perspective on feature influence, describing how sensitive the prediction is to each feature at each time point.

The central contribution of our framework lies in fusing these two attribution modalities. While TreeSHAP reflects global, structural importance, IG captures dynamic, temporal importance. By combining the two, using normalized weighting schemes or multiplicative fusion, we derive a combined feature-importance map that highlights features consistently influential across both modeling paradigms. This fusion works like a reliability filter, highlighting features that both the static and temporal models independently agree on, making it more likely that these signals are real and stable, rather than a model being biased. Overall, this dual-branch interpretability framework not only explains individual model decisions but also provides cross-validated, multiperspective explanations. The resulting stable, trustworthy, and robust attributions are particularly valuable in healthcare and sensor analytics, where interpretability is essential. This approach directly supports the broader objective of trustworthy AI, ensuring that model explanations remain consistent, complementary, and faithful to the underlying decision processes.

## RESULTS

The TADPOLE dataset comprises various types of biomarkers (or measurements), each with varying degrees of importance. The main website (httpxs://tadpole.grand-challenge.org/Data/) recommends focusing on certain key biomarkers initially. Following this suggestion, we selected these key biomarkers along with additional ones, resulting in a total of 44 features. However, due to the presence of numerous null values, we eliminated some features and refined our selection to 22 features. Through rigorous testing, we confirmed that this decision did not negatively impact our model’s performance. This step was crucial to ensure that excluding these features did not lead to losing important information or introducing bias into the model. The final collection of 22 features was chosen based on both their statistical significance and clinical value. These qualities included important clinical assessments including the Alzheimer’s Disease Assessment Scale-Cognitive Subscale (ADAS11, ADAS13), the Mini-Mental State Examination (MMSE), and the Functional Activities Questionnaire (FAQ), among others. We conducted our analysis on these 22 features and identified CDRSB (Clinical Dementia Rating Sum of Boxes) and ADAS13 (Alzheimer’s Disease Assessment Scale) as the top contributors for both Recurrent Neural Networks (RNN) and TreeSHAP interpretations. The dominance of these features made it challenging to observe the contributions of other features. Consequently, we re-ran our analysis excluding CDRSB and ADAS13, focusing on the remaining 20 features. This adjustment led to an 8% decrease in model accuracy but uncovered valuable insights into the importance and interactions of other features.

### Fused Temporal Attributions Reveal Stage-Dependent RAVLT and Clinical Biomarker Importance

In our study, we sought to investigate how different feature sets influence both predictive performance and biological insight in modeling Alzheimer’s disease progression. To this end, we compared two variants of our recurrent neural network (RNN-AD) model: a Base-line Comprehensive Feature Attribution Model, incorporating all 22 features, and a Reduced Feature Attribution Model, derived by ablating two highly dominant features - CDRSB and ADAS13, from the full feature set. These comparisons were performed within our dual-branch interpretability framework, where attributions from the RNN-AD branch were subsequently fused with TreeSHAP-derived attributions from the XG-Boost branch. Both models were trained on the same dataset and evaluated using standard performance metrics, including Macro-Averaged AUC (mAUC) and Balanced Classification Accuracy (BCA). The comprehensive model achieved an mAUC of 0.907 and a BCA of 0.825, while the reduced model achieved an mAUC of 0.881 and a BCA of 0.815. This marginal decrease in performance highlights that even after removing dominant features, the predictive capability of the model remains largely intact. More importantly, this reduction exposes the contribution of less dominant but biologically meaningful features, yielding potentially valuable insights into Alzheimer’s disease progression mechanisms.

For the RNN-AD model, predictions can be configured over a predefined temporal horizon. In this study, we initially forecasted disease progression over 5 years (60 months). Feature attribution was then performed at different prediction intervals using the Integrated Gradients method to capture how feature relevance evolves over time. Since Integrated Gradients assumes the model’s forward function to return a single tensor, we modified the original RNN-AD architecture, originally returning a tuple of tensors (categorical outputs and regression outputs), to separately run simulations for each output type. These IG-derived temporal attributions were later combined with XGBoost-based interaction attributions, forming a consensus fused attribution map.

Thus, two sets of simulations were performed: one focused on categorical classification outputs (CN, MCI, AD) and another on continuous regression outcomes (ADAS11, Ventricles, ICV). In both cases, the model was trained jointly, optimizing a combined objective function integrating categorical cross-entropy and mean absolute error losses. Figure 1 (D) and (E) provide an overview of the biomarker abbreviations and subject demographics for two representative features: RAVLT_immediate (IR) and RAVLT_learning (LR). These distributions underscore the importance of RAVLT measures as potentially significant biomarkers, particularly in relation to diagnostic phase transitions, highlighting their role in disease stage differentiation and longitudinal progression prediction. These RAVLT insights were consistent across both branches of the framework and thus strongly highlighted by the fused attributions.

#### Confirming prior findings and highlighting RAVLT biomarkers in AD progression

##### Baseline Feature Attribution Model

We generated violin plots depicting feature attributions for the cognitively normal (CN) class across four prediction horizons (up to 4 years). Notably, feature attribution patterns remained consistent across time-points. Each diagnostic class, comprising all 22 features along with the baseline diagnostic status (3 additional categorical features), was ranked according to its mean absolute feature attribution scores. These rankings reflect the fused attribution scores, integrating both Tree-SHAP and IG contributions.

For the CN classification, DX: CN and CDRSB emerged as the most influential features, confirming prior findings. Additionally, FAQ, RAVLT_immediate, and ADAS13 ranked among the top 10 contributors. In the MCI classification, DX: MCI was the dominant feature, significantly outweighing all others. For AD classification, DX: AD and FAQ remained the most influential, with ADAS13, RAVLT_perc-forgetting, and DX: AD also contributing substantially.

##### Reduced Feature Attribution Model

We next evaluated a reduced feature set model containing 20 features under the same experimental conditions. Here too, feature attribution patterns remained stable across prediction durations. In CN classification, DX: CN and RAVLT_immediate again ranked highest, followed by FAQ, ABETA_UPENNBIOMK9_04_19_17, and ADAS11 within the top 10. In MCI classification, DX: MCI continued to dominate, and upon excluding this feature, DX: CN emerged as a significant negative predictor of MCI progression. Other relevant features included WholeBrain, FAQ, and MidTemp, although their influence remained secondary. Importantly, these observations were validated across both branches; features highlighted by IG were independently reinforced by TreeSHAP, indicating that the fused attributions represent stable and cross-validated signals.

For AD classification, FAQ was again the most influential predictor, underscoring the pivotal role of baseline MCI status in forecasting AD progression. Additional important contributors included ADAS11, MMSE, and DX: AD.

Overall, ranking attribution scores revealed a strong alignment with previously reported feature hierarchies, reinforcing the robustness of our interpretability framework. However, we also identified additional features, such as RAVLT variants, that represent potentially novel predictors not emphasized in earlier studies. Because these RAVLT features surfaced consistently in both the structural and temporal attribution branches, their emergence in the fused framework suggests they may represent reliable, previously underappreciated biomarkers. Figure 2 (A) summarizes these findings and compares them against prior analyses by Hernandez et. al. ^20^ and El-Sappagh et. al. ^21^.

**Figure 2.**
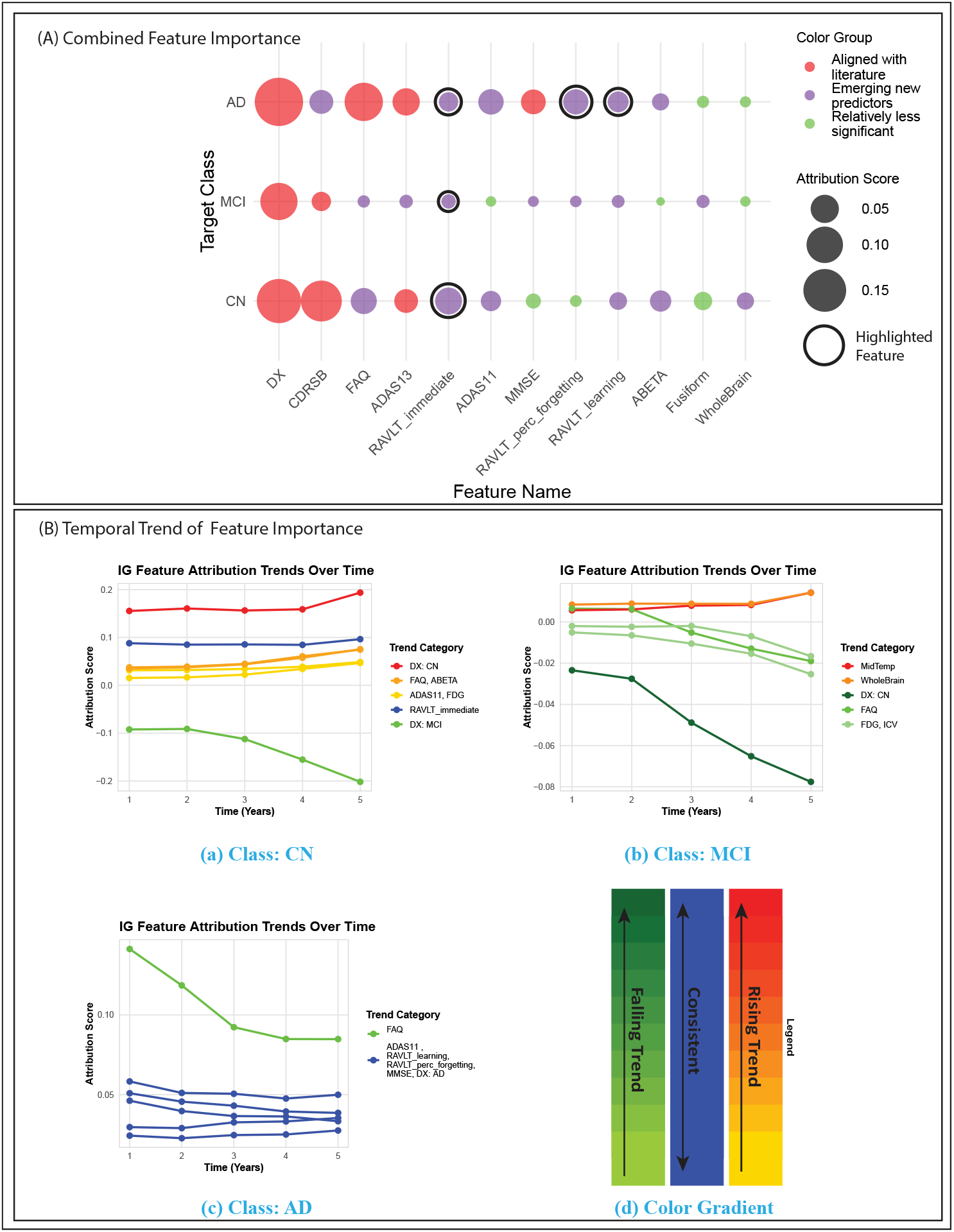
(A) Combined feature importance from fusion framework, (B) Feature attribution trend analysis for 3 stages of Alzheimer’s disease

#### Feature importance values show systematic and smooth transitions over time

We conducted feature attributions across multiple durations to explore potential dependencies and trends in the attributions. Notably, certain features consistently occupy top positions regardless of the duration, while others exhibit varying trends in their attribution scores over time. For the CN class, DX: CN demonstrates a gradual increase in its attribution value as the duration extends. ADAS11, FAQ, FDG and ABETA_UPENNBIOMK9_04_19_17 also show a slow and steady rise in their attribution score. This suggests that these features become increasingly influential in predicting the CN class over longer durations. Conversely, the attribution score for DX: MCI gradually decreases with increasing duration, indicating a reduced likelihood of predicting CN when the baseline diagnosis leans towards MCI. In the case of the MCI class, DX: MCI notably dominates all other features in attribution scores and also demonstrates a skewed rise in its attribution. However, upon discarding this feature, we observe other relevant features with their impacts over time. Features such as DX: CN, FAQ, ICV, FAQ and FDG exhibit slow declining trends in attribution scores, while MidTemp and WholeBrain show a slow but steady rise. However, even though FAQ, ICV, FAQ and FDG show potential for a steady rise in attributions, their values are still negative, which means they can not help predict MCI class. Finally, in the AD class, the significance of FAQ attributions gradually diminishes over time, with its impact becoming negligible towards the later durations. Moreover, DX: MCI always shows high negative attribution throughout the durations, signaling that this feature’s importance does not really help in predicting AD progression as opposed to other relevant features like DX: AD, ADAS11, MMSE, RAVLT_learning and RAVLT_perc_forgetting show consistent influence over time. In summary, the importance or attributions of features indicate how relevant certain features are in predicting a specific diagnosis class. This allows us to derive some biological insights into which features to prioritize when developing diagnostic methods or identifying diagnoses among patients. However, the significance of these features may change over time. Therefore, by analyzing the trends in feature attributions, we can refine and narrow down the most critical features for extraction. A summary of all feature importance trends of specific target classes is depicted in Figure 2 (B). Observing the temporal trends, it is evident that for target class CN, the importance of clinical features shows significant trends, whereas for target class MCI, molecular features such as MidTemp and WholeBrain show visible trends. For the target class AD, we do not see any trend at all. Therefore, we can understand a clear relation between clinical and molecular features and their influences on different prediction target classes.

#### A. SHAP Summary Plot for Target Class CN

### Synergistic and Antagonistic Clinical–Molecular Feature Interactions in Cognitive Decline

The results of this study provide a detailed examination of the interactions between clinical and molecular features across different cognitive states. Utilizing the SHAP library and TreeSHAP methodology, critical insights are uncovered into the relationships that drive Alzheimer’s Disease prediction. The analysis highlights the most interacting features, their unique and shared interactions, and the complex dynamics specific to Alzheimer’s Disease, laying a robust foundation for advancing therapeutic strategies and personalized medicine.

#### Top interacting features across cognitive states

Utilizing the capabilities of the SHAP library enables the generation of a comprehensive summary plot(Figure 3 (A)). This plot illuminates the most prominent features that exhibit substantial interaction (Figure 3 (C)). Interaction is shown in Figure 3 (B). This plot shows the interaction between feature 1 (Hippocampus) and feature 9 (RAVLT_immediate). We can see that the high Hippocampus area (around 8000) and higher value of RAVLT_immediate (30 to 50) are more concerning for the AD class. In the 20-feature model, we observe a greater variety. Some features uniquely exhibit the highest interaction for CN (Cognitively Normal), MCI (Mild Cognitive Impairment), and AD (Alzheimer’s Disease) separately. Within the top seven interacting features, FAQ, ADAS11, RAVLT_immediate, and MMSE consistently emerge across all classes. FAQ, ADAS11, and RAVLT_immediate represent clinical assessments. Some molecular features are also prominent, including Hippocampus, WholeBrain, MidTemp, Fusiform, and Entorhinal. Notably, the MCI class is distinguished by unique molecular features such as Whole-Brain, MidTemp, and Fusiform, while the AD class is characterized by Entorhinal and the CN class by RAVLT_learning.

**Figure 3.**
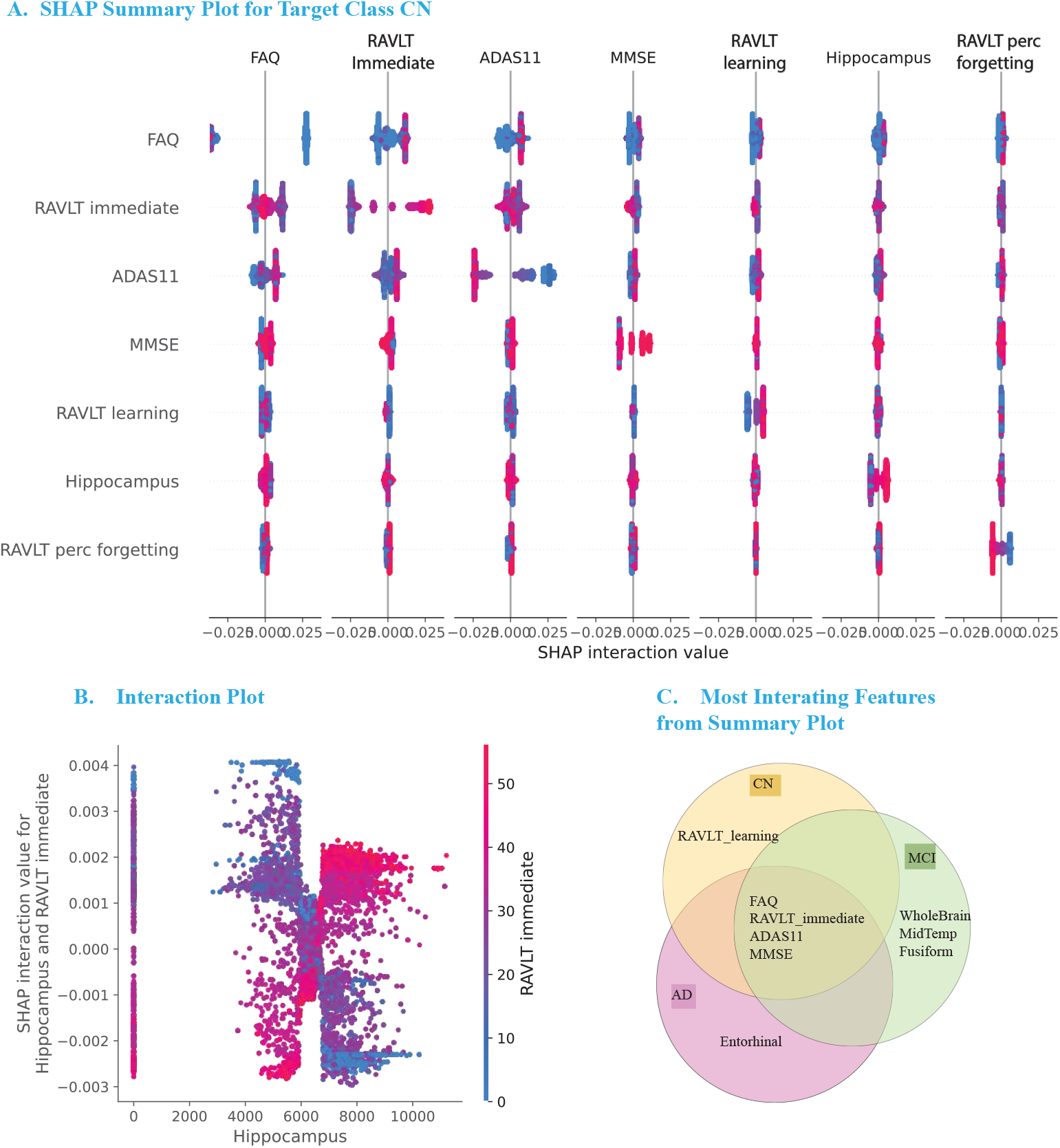
(A) Comprehensive summary plot for target stage CN (Cognitive Normal), (B) Interaction plot of Hippocampus and RAVLT_immediate and (C) Venn diagram of most interacting features across all 3 target stages

#### Clinical and Molecular features reveal complex interaction networks driving Alzheimer’s prediction

We can divide all features into two categories: clinical and molecular (Figure 4 (A)). All cognitive tests fall under the clinical category, while features such as brain structural integrity, proteins and genes are classified as molecular or biological features. For a comprehensive understanding of interactions and their ranking, we have generated interaction value matrices for all classes(supplement). These matrices allow for a clear visualization of interactions and their relative importance. Subsequently, the top 12 features extracted from these matrices are presented in a Venn diagram (Figure 4 (A)) for further analysis. In the top twelve features, all clinical attributes remain consistent across classes, albeit with varying ranks. An intriguing observation emerges in the 20-feature model, where almost half of the top features are clinical across all classes. Notably, molecular or biological features exhibit a diverse range both in terms of rank and specific attributes. Common to all three classes are the features of Entorhinal and Whole-Brain, each with distinct rankings. Additionally, the CN and MCI classes share the feature ICV, while the CN and AD classes share Hippocampus and AV45, and the MCI and AD classes share MidTemp and Fusiform.

**Figure 4.**
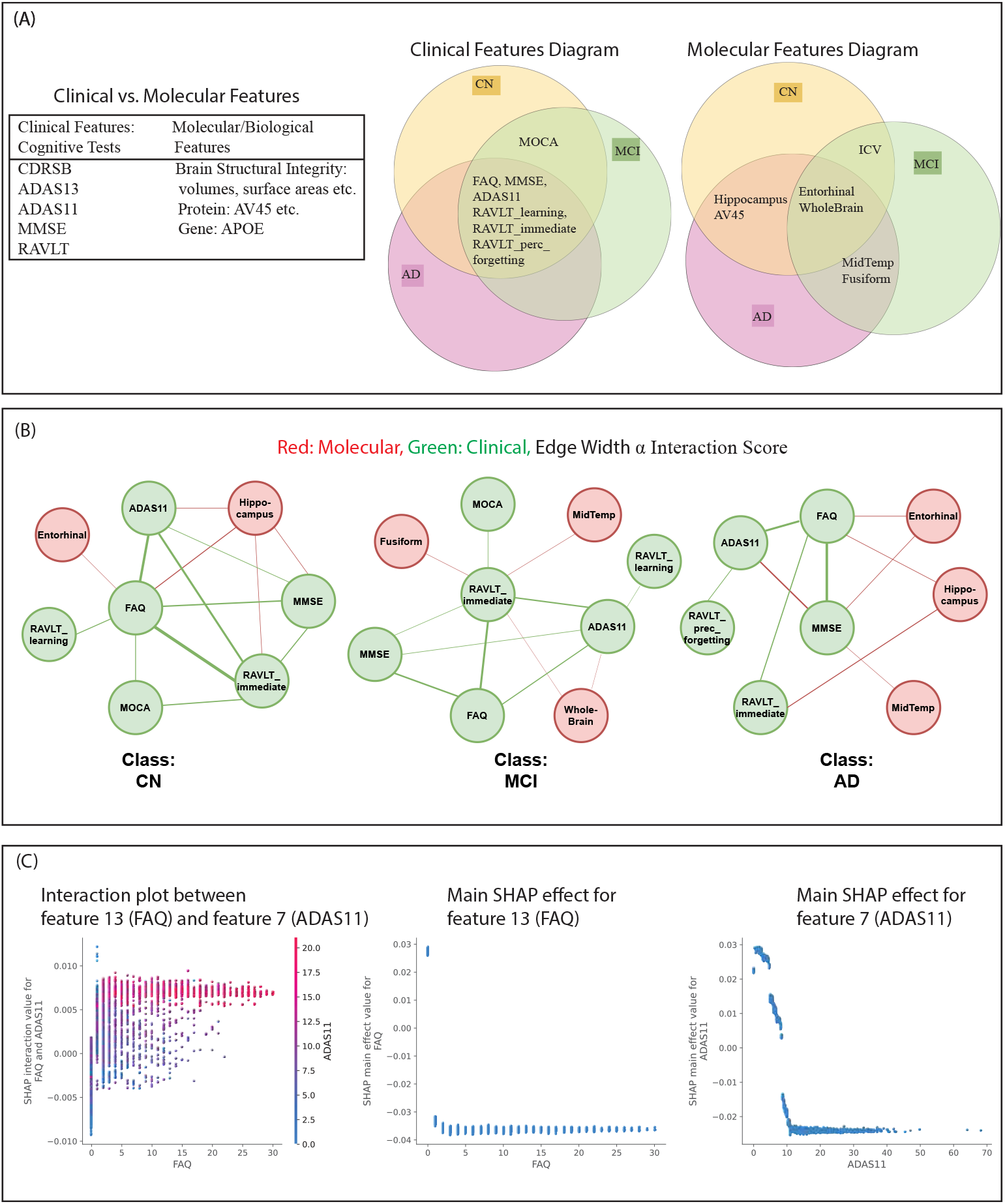
(A)Clinical and molecular feature distinction with their respective influence over 3 stages of prediction, (B) A graphical representation of interactions over 3 stages of prediction (C) Example relation between FAQ and ADAS11 with the help of interaction plot

From interaction value matrices, we have generated a graph by choosing some interesting interaction pairs (Figure 4 (B)). Each node in the graph represents a feature. Nodes marked in red are molecular features, while the others are clinical features. The weight of the edges represents the interaction values of those features from the matrix. For cognitively normal (CN) individuals, fourteen notable interactions are observed, comprising clinical-clinical and clinical-molecular pairings, with clinical variables predominating. FAQ emerges as the most frequently occurring variable, highlighting its central role in these interactions. The molecular feature Hippocampus shows significant interactions with clinical attributes such as MMSE, FAQ, RAVLT_immediate, and ADAS11. In Mild Cognitive Impairment (MCI), the graph illustrates diverse interactions among twelve ranked pairs, mainly clinical-clinical. RAVLT_immediate is the primary interacting feature, centrally positioned with connections to clinical features like MMSE, FAQ, ADAS11, MOCA, and molecular features such as WholeBrain, Fusiform, and MidTemp. In Alzheimer’s disease (AD), the graph displays a rich set of interactions, with clinical-clinical pairings predominating. FAQ, MMSE, and ADAS11 are central clinical features, indicating their significant roles in the interaction network. FAQ demonstrates extensive connectivity, emphasizing its importance across various pairs. The Hippocampus also exhibits significant interactions with clinical attributes such as MMSE, FAQ, and RAVLT_immediate. An intriguing discovery regarding the relationship between FAQ and ADAS11 in the context of CN is revealed when the interaction plot and SHAP plot of these two features are plotted in Figure 4 (C). The individual SHAP plot shows that higher FAQ and ADAS11 values negatively impact the model. In contrast, the interaction plot reveals that elevated FAQ and ADAS11 levels increase the likelihood of detecting CN. This effect is not evident for MCI or AD. Thus, identical feature pairs interact differently across different classes.

### A Unified Interaction–Trend Framework for AD Biomarker Interpretation

Across the CN, MCI, and AD groups, the interaction– trend quadrant maps in Figure 5 reveal distinct and stage-specific patterns in how clinical and molecular features contribute to disease characterization. Clinical assessments such as ADAS11, MMSE, and FAQ consistently occupy the high-interaction quadrants, indicating that they influence the prediction not only directly but also through strong pairwise interactions with other features. Their temporal slopes reveal additional nuances: in CN and MCI, these cognitive scores either slightly increase or remain stable, indicating that they become more relevant as people move away from normal aging. However, in AD, these same measures shift toward declining trends, reflecting ceiling effects and reduced discriminative power in later stages.

**Figure 5.**
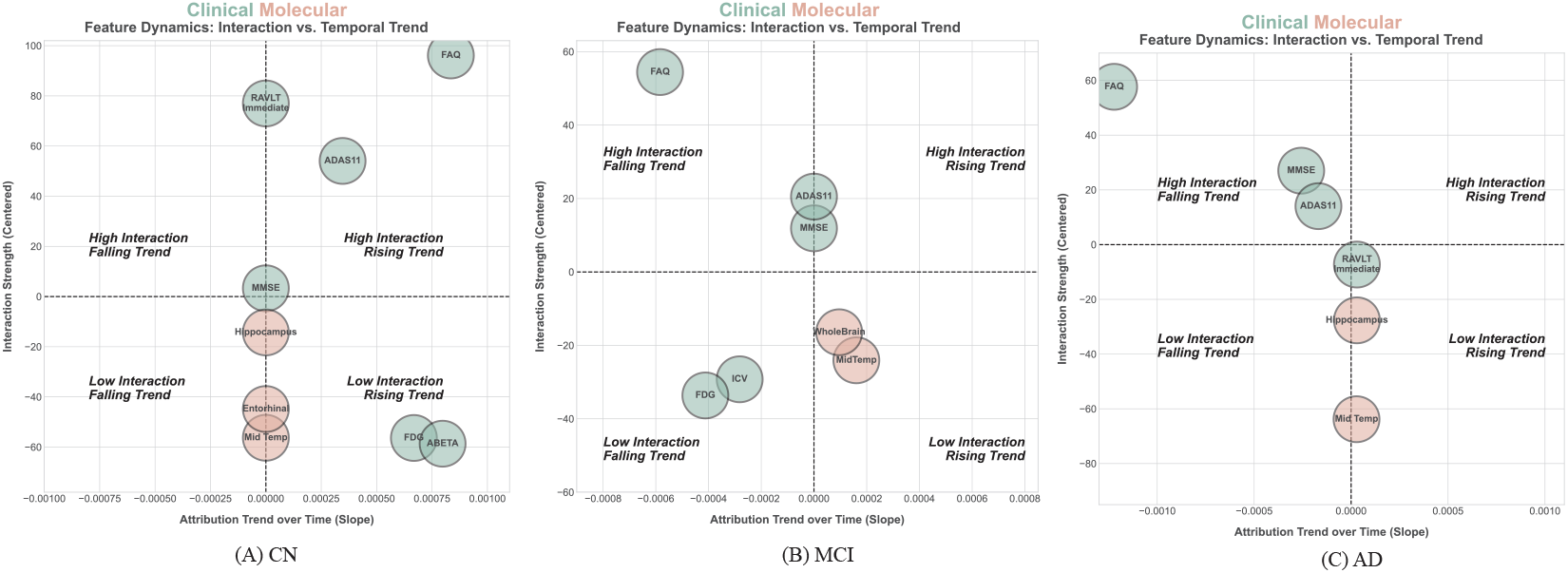
Unified analysis of feature interaction and feature importance trend

In contrast, structural MRI features such as Hip-pocampus, Entorhinal, Mid Temp, and WholeBrain predominantly fall into the low-interaction quadrants. These features show steadily decreasing importance over time. This pattern fits the biological course of Alzheimer’s, where brain degeneration starts early and then saturates, making these markers less useful as the disease progresses. Notably, metabolic markers such as FDG and molecular indicators like ABETA show stage-dependent shifts: moderate interaction strength in CN, declining importance in MCI, and re-emerging relevance in AD. This suggests that non-linear trajectories are potentially linked to the brain’s compensatory responses or different mechanisms active at each stage. Together, these findings highlight that no single biomarker remains uniformly informative across all disease stages. Instead, clinical scores gain importance through interactions in early disease stages, while imaging and molecular markers are most sensitive early on but lose value later. This interaction–trend framework provides a powerful analogy for identifying stage-adaptive biomarkers and for constructing dynamic, multimodal models that reflect real biological progression.

## DISCUSSION

### Feature Importance Dynamics as a Basis for Improved Clinical Trials and Therapeutic Strategies

It is important to mention that a temporal model like RNN-AD does not consider diagnosis as a feature. However, feature attributions can be found for the categorical features (DX:CN, DX:MCI, DX:AD). Intuitively, for a temporal model like RNN-AD, where future timeline predictions are somewhat influenced by the predictions of previous timelines, attributions of earlier timelines are more significant than the later ones. These findings shed light on the evolving nature of feature importance in predicting ‘AD’ progression over different time intervals. While some features maintain their relevance consistently, others demonstrate varying trends in attribution scores, reflecting the complex interplay between biomarkers and disease progression. Understanding these temporal dynamics can inform the development of more accurate and robust predictive models for tracking disease progression and guiding clinical interventions in patients with Alzheimer’s disease. By identifying the temporal patterns of feature importance, clinicians and researchers can prioritize the monitoring of specific biomarkers and clinical assessments at different stages of disease progression. This tailored approach enables early detection of cognitive decline, facilitates personalized treatment strategies and trial design, and ultimately enhances patient care and management. Moreover, understanding the temporal dynamics of feature attributions can also guide the selection of features for model refinement and optimization. Features that exhibit consistent importance across durations may warrant further investigation for their predictive value and underlying biological significance. Conversely, features with fluctuating attributions may indicate dynamic changes in disease pathology or response to treatment, highlighting the need for adaptive modeling approaches.

### Synergistic and Antagonistic Clinical–Molecular Relationships Inspire Improved Feature Engineering

The comprehensive analysis conducted in this study offers substantial insights into the interactions between clinical and molecular features across different cognitive states. By employing the SHAP library and TreeSHAP methodology, critical relationships driving Alzheimer’s Disease prediction have been identified. This analysis underscores the importance of specific interacting features, highlighting both their unique and shared interactions, as well as the complex dynamics specific to Alzheimer’s Disease. These findings provide a robust foundation for advancing therapeutic strategies and personalized medicine. The study reveals that a subset of features consistently exhibits significant interactions across various cognitive states, including Cognitively Normal (CN), Mild Cognitive Impairment (MCI), and Alzheimer’s Disease (AD). For instance, in the 20-feature model, key clinical features such as FAQ, ADAS11, RAVLT_immediate, and MMSE consistently emerge across all classes. The 22-feature model introduces additional clinical features like CDRSB and ADAS13, which further contribute to the understanding of feature interactions. In cognitively normal individuals, the analysis identifies fourteen notable interactions predominantly involving clinical-clinical pairings, with some clinical-molecular combinations. FAQ emerges as a central clinical feature, frequently interacting with other variables, while the molecular feature Hippocampus shows significant interactions with clinical attributes like MMSE, FAQ, RAVLT_immediate, and ADAS11. This pattern is consistent in the 22-feature model, where clinical features remain predominant, and CDRSB becomes the top interacting feature. For individuals with Mild Cognitive Impairment, the study highlights a diverse array of interactions, predominantly clinical-clinical, with RAVLT_immediate as a primary interacting feature. This feature interacts with multiple clinical and molecular attributes, such as WholeBrain, Fusiform, and MidTemp. In the 22-feature model, a shift in interaction dynamics is observed, with MMSE, FAQ, and CDRSB emerging as key interacting features. The presence of molecular features interacting with clinical ones is reduced, underscoring the variability in feature interactions between the models. In Alzheimer’s Disease, the analysis reveals rich interactions primarily involving clinical-clinical pairings, with notable clinical-molecular interactions. FAQ, MMSE, and ADAS11 are central clinical features, with Hippocampus also showing significant interactions with clinical attributes. The 22-feature model introduces CDRSB and ADAS13 as top interacting features, adding complexity to the interaction network. These findings enhance the understanding of the interplay between different clinical and molecular features in Alzheimer’s Disease. Overall, this study provides valuable insights into the complex interactions between clinical and molecular features in Alzheimer’s Disease prediction. The identified interactions and their dynamics offer a foundation for future research and therapeutic advancements, emphasizing the need for personalized approaches in the treatment and management of Alzheimer’s Disease.

## REFERENCES

1. Blennow K, de Leon MJ, Zetterberg H. Alzheimer’s disease. The Lancet. 2006;368(9533):387–403.

2. Guerreiro R, Bras J. The age factor in Alzheimer’s disease. Genome medicine. 2015;7:1–3.

3. Reeve A, Simcox E, Turnbull D. Ageing and Parkinson’s disease: why is advancing age the biggest risk factor? Ageing research reviews. 2014;14:19–30.

4. Piemontese L. New approaches for prevention and treatment of Alzheimer’s disease: a fascinating challenge. Neural regeneration research. 2017;12(3):405–6.

5. Maitra U, Stephen C, Ciesla LM. Drug discovery from natural products–Old problems and novel solutions for the treatment of neurodegenerative diseases. Journal of pharmaceutical and biomedical analysis. 2022;210:114553.

6. Rasmussen J, Langerman H. Alzheimer’s disease–why we need early diagnosis. Degenerative neurological and neuromuscular disease. 2019:123–30.

7. Prescott JW. Quantitative imaging biomarkers: the application of advanced image processing and analysis to clinical and preclinical decision making. Journal of digital imaging. 2013;26:97–108.

8. Hua X, Leow AD, Parikshak N, Lee S, Chiang MC, Toga AW, et al. Tensor-based morphometry as a neuroimaging biomarker for Alzheimer’s disease: an MRI study of 676 AD, MCI, and normal subjects. Neuroimage. 2008;43(3):458–69.

9. Ebrahimighahnavieh MA, Luo S, Chiong R. Deep learning to detect Alzheimer’s disease from neuroimaging: A systematic literature review. Computer methods and programs in biomedicine. 2020;187:105242.

10. Wen J, Thibeau-Sutre E, Diaz-Melo M, Samper-González J, Routier A, Bottani S, et al. Convolutional neural networks for classification of Alzheimer’s disease: Overview and reproducible evaluation. Medical image analysis. 2020;63:101694.

11. Marinescu RV, Oxtoby NP, Young AL, Bron EE, Toga AW, Weiner MW, et al. TADPOLE challenge: prediction of longitudinal evolution in Alzheimer’s disease. arXiv preprint arXiv:180503909. 2018.

12. Bron EE, Smits M, Van Der Flier WM, Vrenken H, Barkhof F, Scheltens P, et al. Standardized evaluation of algorithms for computer-aided diagnosis of dementia based on structural MRI: the CADDementia challenge. NeuroImage. 2015;111:562–79.

13. Allen GI, Amoroso N, Anghel C, Balagurusamy V, Bare CJ, Beaton D, et al. Crowdsourced estimation of cognitive decline and resilience in Alzheimer’s disease. Alzheimer’s & Dementia. 2016;12(6):645–53.

14. Marinescu RV, Bron EE, Oxtoby NP, Young AL, Toga AW, Weiner MW, et al. Predicting Alzheimer’s disease progression: results from the TADPOLE challenge: neuroimaging: neuroimaging predictors of cognitive decline. Alzheimer’s & Dementia. 2020;16:e039538.

15. Sauty B, Maheux E, Durrleman S. Feature Selection to Forecast Cognitive Decline Using Multimodal Alzheimer’s Disease Models. 2023.

16. Tavakoli HM, Xie T, Shi J, Hadzikadic M, Ge Y. Predicting neural deterioration in patients with alzheimer’s disease using a convolutional neural network. In: 2020 IEEE International Conference on Bioinformatics and Biomedicine (BIBM). IEEE; 2020. p. 1951–8.

17. Jung W, Jun E, Suk HI, Initiative ADN, et al. Deep recurrent model for individualized prediction of Alzheimer’s disease progression. NeuroImage. 2021;237:118143.

18. Viswan V, Shaffi N, Mahmud M, Subramanian K, Hajamohideen F. Explainable artificial intelligence in Alzheimer’s disease classification: A systematic review. Cognitive Computation. 2024;16(1):1–44.

19. Lundberg SM, Lee SI. A unified approach to interpreting model predictions. Advances in neural information processing systems. 2017;30.

20. Hernandez M, Ramon-Julvez U, Ferraz F, with the ADNI Consortium. Explainable AI toward understanding the performance of the top three TADPOLE Challenge methods in the forecast of Alzheimer’s disease diagnosis. PloS one. 2022;17(5):e0264695.

21. El-Sappagh S, Alonso JM, Islam SR, Sultan AM, Kwak KS. A multilayer multimodal detection and prediction model based on explainable artificial intelligence for Alzheimer’s disease. Scientific reports. 2021;11(1):2660.

22. Moradi E, Hallikainen I, Hänninen T, Tohka J, Initiative ADN, et al. Rey’s Auditory Verbal Learning Test scores can be predicted from whole brain MRI in Alzheimer’s disease. NeuroImage: Clinical. 2017;13:415–27.

23. Kueper JK, Speechley M, Montero-Odasso M. The Alzheimer’s disease assessment scale–cognitive subscale (ADAS-Cog): modifications and responsiveness in predementia populations. a narrative review. Journal of Alzheimer’s Disease. 2018;63(2):423–44.

24. Lansdall CJ, McDougall F, Butler L, Delmar P, Pross N, Qin S, et al. Establishing clinically meaningful change on outcome assessments frequently used in trials of mild cognitive impairment due to Alzheimer’s disease. The journal of prevention of Alzheimer’s disease. 2023;10(1):9–18.

25. Chen M. Minimalrnn: Toward more interpretable and trainable recurrent neural networks. arXiv preprint arXiv:171106788. 2017.

26. Chen T, Guestrin C. Xgboost: A scalable tree boosting system. In: Proceedings of the 22nd acm sigkdd international conference on knowledge discovery and data mining; 2016. p. 785–94.

27. Nguyen M, He T, An L, Alexander DC, Feng J, Yeo BT, et al. Predicting Alzheimer’s disease progression using deep recurrent neural networks. NeuroImage. 2020;222:117203.

28. Sundararajan M, Taly A, Yan Q. Axiomatic attribution for deep networks. In: International conference on machine learning. PMLR; 2017. p. 3319–28.

29. Ribeiro MT, Singh S, Guestrin C. “Why should i trust you?” Explaining the predictions of any classifier. In: Proceedings of the 22nd ACM SIGKDD international conference on knowledge discovery and data mining; 2016. p. 1135–44.

30. Shrikumar A, Greenside P, Kundaje A. Learning important features through propagating activation differences. In: International conference on machine learning. PMLR; 2017. p. 3145–53.

31. Montavon G, Binder A, Lapuschkin S, Samek W, Müller KR. Layer-wise relevance propagation: an overview. Explainable AI: interpreting, explaining and visualizing deep learning. 2019:193–209.

32. Wang D, Honnorat N, Fox PT, Ritter K, Eickhoff SB, Seshadri S, et al. Deep neural network heatmaps capture Alzheimer’s disease patterns reported in a large meta-analysis of neuroimaging studies. Neuroimage. 2023;269:119929.

33. Čík I, Rasamoelina AD, Mach M, Sinčák P. Explaining deep neural network using layer-wise relevance propagation and integrated gradients. In: 2021 IEEE 19th world symposium on applied machine intelligence and informatics (SAMI). IEEE; 2021. p. 000381–6.

34. Dhamdhere K, Sundararajan M, Yan Q. How important is a neuron? arXiv preprint arXiv:180512233. 2018.

35. Lundberg SM, Erion GG, Lee SI. Consistent individualized feature attribution for tree ensembles. arXiv preprint arXiv:180203888. 2018.

