## Supplementary for "Modeling Temporal Dependencies and Feature Interactions Reveal Novel Clinical and Molecular Insights into Alzheimer’s Disease Progression"

*December 25, 2024*

#### 1 Disease Biomarkers

Quantitative biomarkers are medical measurements that can indicate a disease. The TADPOLE dataset contains the following fields.

1. Main Cognitive Tests:
   - CDR Sum of Boxes (CDRSB)
   - ADAS11
   - ADAS13
   - MMSE
   - RAVLT
   - Moca
   - Ecog
2. MRI ROIs (Freesurfer) - measures of brain structural integrity
   - volumes
   - cortical thicknesses
   - surface areas
3. FDG PET ROI averages - measure cell metabolism, where cells affected by AD show reduced metabolism
4. AV45 PET ROI averages - measures amyloid-beta load in the brain, where amyloid-beta is a protein that misfolds (i.e. its 3D structure is not properly constructed), which then leads to AD
5. AV1451 PET ROI averages - measures tau load in the brain, where tau is another protein which, when abnormal, damages neurons and thus leads to AD
6. DTI ROI measures - measures microstructural parameters related to cells and axons (cell radial diffusivity, axonal diffusivity, etc ... )
   - Mean diffusivity
   - Axial diffusivity
   - Radial diffusivity
7. CSF biomarkers - amyloid and tau levels in the cerebrospinal fluid (CSF), as opposed to the cerebral cortex
8. Others: APOE status, demographic information, Diagnosis: either cognitively normal (CN), mild cognitive impairment (MCI), or Alzheimer's disease (AD).

### 2 dataset

| | | |
|---|---|---|
| D1 | Standard Training Set | Contains regional MRI (volumes, cortical thickness, surface area), PET measures (ROI SUVR values: FDG, AV45, AV1451), DTI (ROI summary: regional means of standard indices) and CSF measures (Elecsys analysis: Amyloid-beta, Tau and P-Tau) |
| D2 | Standard Prediction Set | Contains all currently available longitudinal data for prospective ADNI-3 subjects that are rollovers from earlier ADNI studies. Such subjects are active (PTSTATUS==1), with ADNI-2 visits (Phase==ADNI2), and screening was performed (RGSTATUS==’1’) |
| D3 | Cross Sectional Prediction Set | Includes the same set of patients that appeared in D2, with only the final visit and a few data columns in order to mimic screening data for a clinical trial. In this case, the available information is typically limited to demographics, cognitive test scores, and structural MRI (derived brain volumes). |
| D4 | Test Set | ADNI-3 test data containing rollover individuals. This is acquired after the challenge submission deadline. Evaluation on the forecasts are done with this dataset according to the challenge metrics. |

Table 1: Dataset summary

| **Bio markers** | **Corresponding Features Number** |
|---|---|
| Ventricles | 0 |
| Hippocampus | 1 |
| WholeBrain | 2 |
| Entorhinal | 3 |
| Fusiform | 4 |
| MidTemp | 5 |
| ICV | 6 |
| ADAS11 | 7 |
| MMSE | 8 |
| RAVLT_immediate | 9 |
| RAVLT_learning | 10 |
| RAVLT_forgetting | 11 |
| RAVLT_perc_forgetting | 12 |
| FAQ | 13 |
| MOCA | 14 |
| AV45 | 15 |
| FDG | 16 |
| ABETA_UPENNBIOMK9_04_19_17 | 17 |
| TAU_UPENNBIOMK9_04_19_17 | 18 |
| PTAU_UPENNBIOMK9_04_19_17 | 19 |

Table 2: Biomarkers and their feature number

### 3 Figures

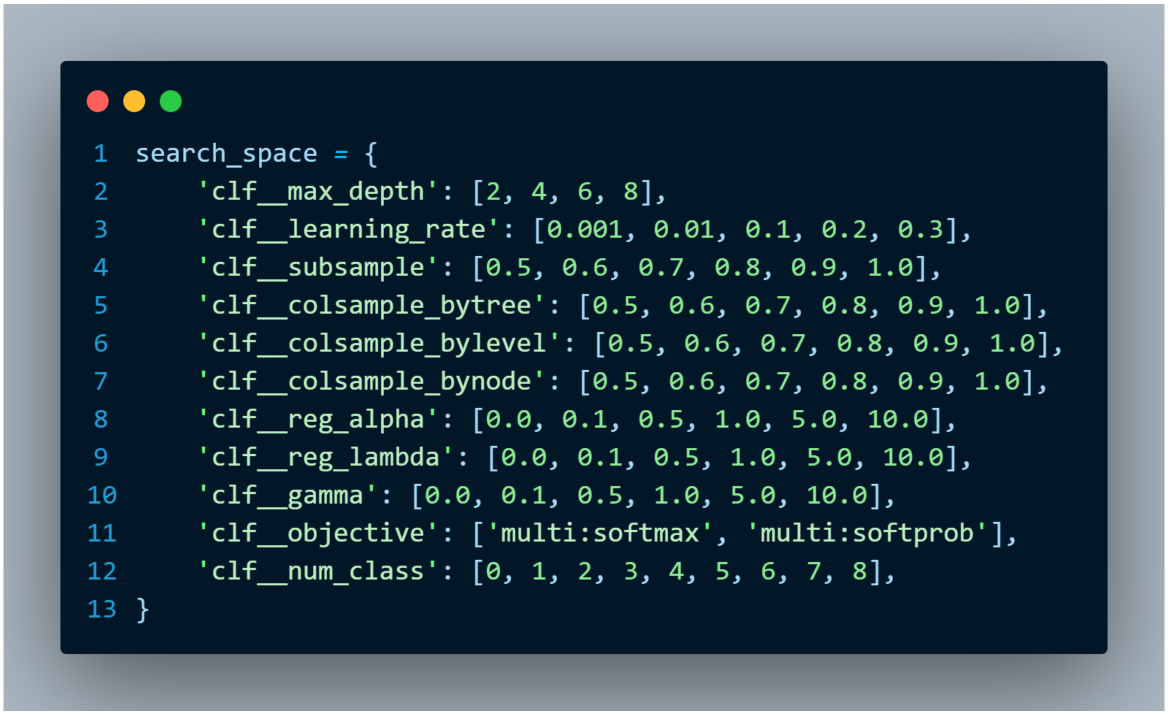

```
search_space = {
    'clf__max_depth': [2, 4, 6, 8],
    'clf__learning_rate': [0.001, 0.01, 0.1, 0.2, 0.3],
    'clf__subsample': [0.5, 0.6, 0.7, 0.8, 0.9, 1.0],
    'clf__colsample_bytree': [0.5, 0.6, 0.7, 0.8, 0.9, 1.0],
    'clf__colsample_bylevel': [0.5, 0.6, 0.7, 0.8, 0.9, 1.0],
    'clf__colsample_bynode': [0.5, 0.6, 0.7, 0.8, 0.9, 1.0],
    'clf__reg_alpha': [0.0, 0.1, 0.5, 1.0, 5.0, 10.0],
    'clf__reg_lambda': [0.0, 0.1, 0.5, 1.0, 5.0, 10.0],
    'clf__gamma': [0.0, 0.1, 0.5, 1.0, 5.0, 10.0],
    'clf__objective': ['multi:softmax', 'multi:softprob'],
    'clf__num_class': [0, 1, 2, 3, 4, 5, 6, 7, 8],
}
```

Figure 1: XGBoost search space code

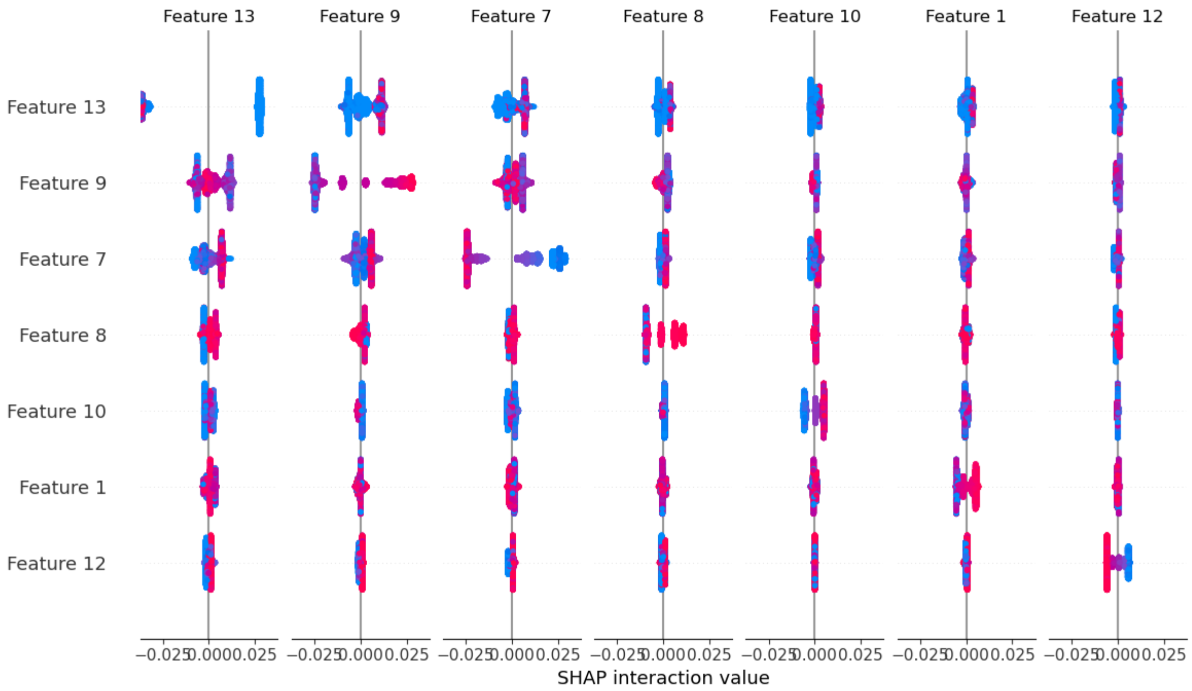


Figure 2: SHAP summary plot for CN [Table: 2]

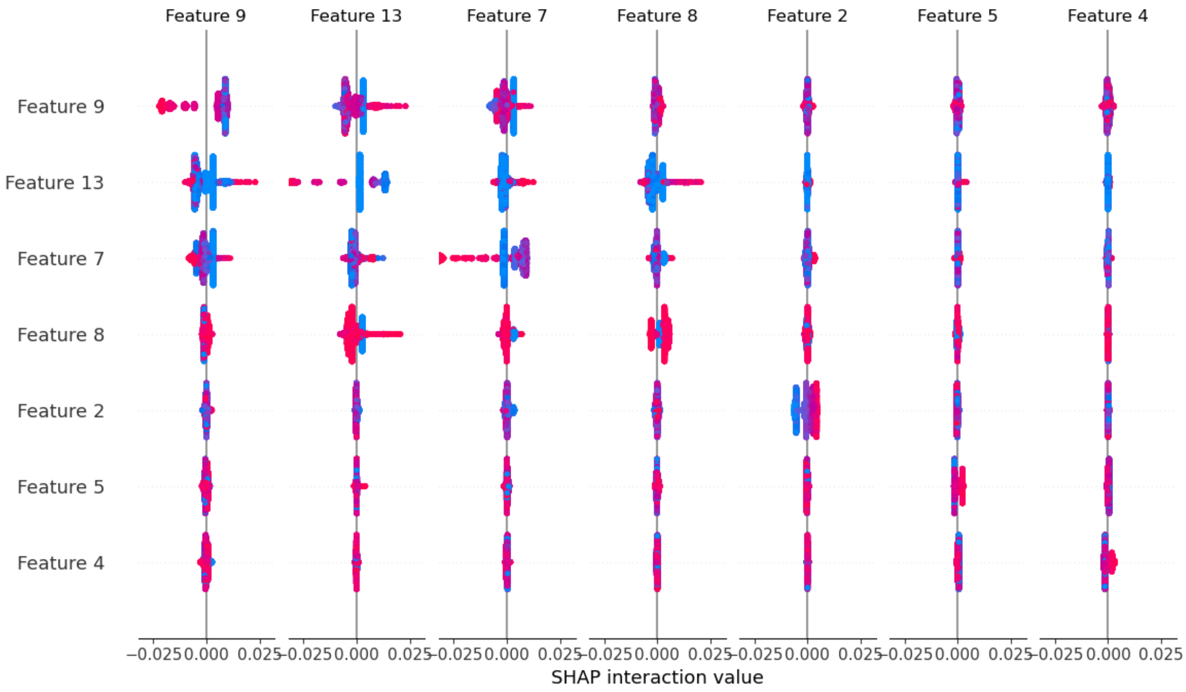


Figure 3: SHAP summary plot for MCI [Table: 2]

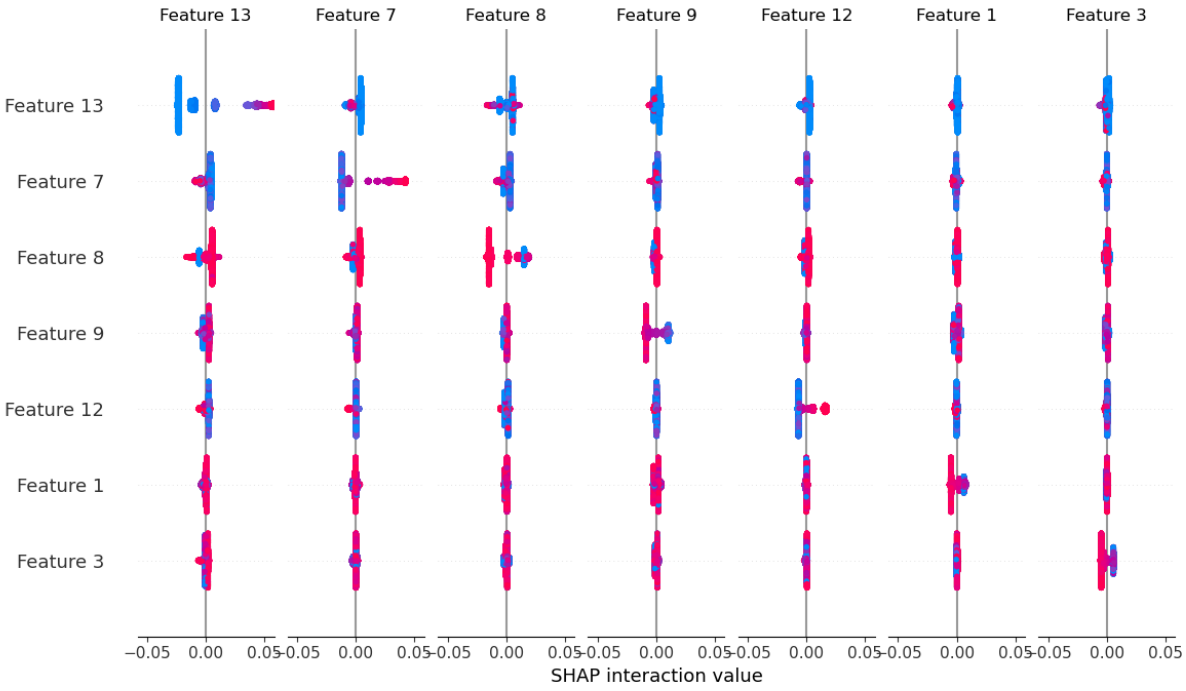


Figure 4: SHAP summary plot for AD [Table: 2]
